# Cholinergic signalling in the forebrain controls microglial phenotype and responses to systemic inflammation

**DOI:** 10.1101/2021.01.18.427123

**Authors:** Arshed Nazmi, Eadaoin W. Griffin, Robert H. Field, Sean Doyle, Edel Hennessy, Martin O’Donnell, Aisling Rehill, Anthony McCarthy, Daire Healy, Michelle M. Doran, John P. Lowry, Colm Cunningham

## Abstract

Loss of basal forebrain cholinergic projections occurs in Alzheimer’s disease, frontotemporal dementia and in aging. Moreover, nicotinic stimulation is anti-inflammatory in macrophages and microglia but how loss of basal forebrain acetylcholine impacts on microglial phenotype is poorly understood. Here we hypothesized that endogenous ACh maintains homeostatic microglial phenotype and that neurodegeneration-evoked loss of ACh tone, triggers microglial activation. Using the specific immunotoxin, mu-p75^NTR^-saporin, we performed partial lesions of the basal forebrain cholinergic nuclei, medial septum and ventral diagonal band. We examined microglial phenotype in the hippocampus, the major projection area for these nuclei, using bulk RNA preparations, Flow cytometry-sorted microglial cells, immunohistochemistry and ELISA to examine responses to cholinergic withdrawal and acute responses to subsequent systemic inflammation with LPS. Basal forebrain cholinergic degeneration elicited lasting activation of microglia in the hippocampus, showing suppression of *Sall1* and persistent elevation of *Trem2, Clec7a, Itgax* and complement genes proportionate to *Chat* loss. These primed microglia showed exaggerated IL-1β responses to systemic LPS challenge. In normal animals LPS evoked acute increases in extracellular choline, a proxy for ACh release, and this response was lost in lesioned animals. Restoration of basal cholinergic signalling via serial treatments with the nicotinic agonist PNU282,987 resulted in reversion to the homeostatic microglial phenotype and prevented exaggerated responses to acute systemic inflammation. The data indicate that neurodegeneration-evoked loss of cholinergic tone, triggers microglial activation via impaired microglial nicotinic signalling and leaves these microglia more vulnerable to secondary inflammatory insults. The data have implications for neuroinflammation during aging and neurodegeneration and for responses to sepsis and systemic inflammation.

## Introduction

In Alzheimer’s disease (AD), frontotemporal dementia (FTD) and aging there is significant loss of basal forebrain cholinergic projections to the hippocampus and cortex (Davies & Maloney, 1976; Whitehouse *et al*., 1982; Nyakas *et al*., 2011). This reduces Acetylcholine (ACh) neuromodulatory influence on these brain structures and use of acetylcholinesterase inhibitors, to enhance ACh levels, remains a first line treatment for Alzheimer’s disease. ACh has also been shown to exert anti-inflammatory actions on macrophages and microglia but the impact of degeneration in the cholinergic system for microglial phenotype is poorly understood.

Early evidence for anti-inflammatory influence of ACh, came from vaogotomy experiments showing that vagal nerve-released ACh exerts anti-inflammatory control over TNF-α and IL-1β secretion from splenic macrophages. This effect was mimicked by agonists of the α7 nicotinic receptor (α_7_nAChR) (Borovikova *et al*., 2000; Pavlov *et al*., 2003; Wang *et al*., 2003). This peripheral pathway is also relevant to neuroinflammation: mice subjected to sleep deprivation, or to tibial fracture coupled with endotoxemia or ischemia, revealed anti-inflammatory effects of α_7_nAChR agonist, via modulation of peripheral monocytes (Terrando *et al*., 2015; Xue *et al*., 2019) and/or macrophage infiltration (Han *et al*., 2014), with downstream effects on neuroinflammation.

*In vitro* studies in microglia, showed that nicotine and ACh suppressed LPS-induced TNF-α and IL-1β in both rat and mouse microglia (Shytle *et al*., 2004; De Simone *et al*., 2005), while the acetylcholine esterase inhibitor, galantamine, suppressed HIV-1 gp120-mediated microglial TNF-α secretion from microglia and this was blunted by the nicotinic antagonist α-bungarotoxin (Giunta *et al*., 2004). The nicotinic agonist DMXBA also increased microglial phagocytosis of Aβ *in vitro* (Takata *et al*., 2018).

Basal forebrain cholinergic neurons release ACh in multiple brain areas with abundant nAChRs (Fabian-Fine *et al*., 2001) and with high densities of microglia and astrocytes (Jinno *et al*., 2007) but *in vivo* evidence for a direct anti-inflammatory effect of basal ACh on microglia is not clear. Peripheral injections and chronic infusions of nicotine have opposite effects on the microglial marker Iba1 in the nucleus accumbens of P32 and P90 rats (Linker *et al*., 2020). Chronic nicotine treatment reduced the loss of dopaminergic neurons in the MPTP model of Parkinson’s disease (Park *et al*., 2007; Liu *et al*., 2012a) but it is not clear whether this was mediated by direct effects on microglia or on dopaminergic neurons (Janson *et al*., 1992).

Thus, despite the plausibility of a CNS cholinergic anti-inflammatory pathway, there is limited *in vivo* evidence that microglia are directly influenced by endogenous acetylcholine. Moreover, despite the enormous interest in microglial activation in AD (Hickman *et al*., 2018), it is striking that we do not have clear information on how the loss of cholinergic tone to the forebrain, which clearly occurs in AD (Whitehouse *et al*., 1981; Hampel *et al*., 2018), affects the phenotype of microglial cells in the hippocampus and cortex. Here we hypothesized that endogenous levels of ACh are necessary to maintain microglial homeostasis. In addition to the adoption of a specific activated transcriptional profile, these microglia were predicted to be primed by loss of cholinergic tone to show exaggerated responses to subsequent inflammatory stimulation. Such ‘primed’ responses have been shown in other models of neurodegeneration (Holtman *et al*., 2015), but understanding of the mechanisms by which microglial become primed remains limited. We propose that the loss of ACh tone is among the triggers for loss of microglial homeostasis, with resulting priming. In the current study we have performed partial lesions of the basal forebrain cholinergic nucleus and examined the phenotype of isolated microglial cells from the projection areas for these cholinergic nuclei. We demonstrate a loss of homeostatic phenotype and subsequent LPS-induced switching to additional microglial states and demonstrate the profound influence of cholinergic signaling on the control of pro-inflammatory microglial responses.

## Methods

### Surgery and animal treatments

Female C57BL/6 mice (HarlanOlac, UK) were housed in groups of five at 21°C with a 12:12 h light-dark cycle with food and water ad libitum. Mice were anesthetized intraperitoneally (i.p.) with Avertin (2,2,2-tribromoethanol; Sigma) and positioned in a stereotaxic frame (Kopf Instruments). Two (i.e. bilateral) 1 µl injections of p75^NTR^-Saporin (Advanced Targeting Systems) at a concentration of either 0.2 or 0.6 µg/µl (total dosage, 0.4 µg or 1.2 µg respectively) were made into the lateral ventricles (i.c.v) using a pulled glass micropipette [coordinates (mm) from bregma: anteroposterior, −0.22; lateral, ±1.0; depth, −1.9]. Control animals were injected with sterile PBS. Following injection, the needle remained in place for 5 min to minimize reflux. Animals recovered fully from surgical procedures (≥ 14d) before giving other treatments. Animals were euthanized at various time points post-lesion to assess temporal evolution of microglial changes and a number of additional animals were injected (i.p) with either LPS (100µg/kg) or saline at 35-40 days post-lesion in order to examine microglial responses to acute systemic inflammation. For PNU experiments a cohort of female C57BL/6 mice were injected with either p75-Saporin (0.6µg bilaterally) or PBS i.c.v. Animals were then administered PNU-282,987 (i.p.) in one of two doses, 10mg/kg or 20mg/kg, or a vehicle control. Animals were perfused at 38 days and brain and blood samples harvested. For flow cytometry and cell sorting experiments, a cohort of female mice was injected with 1.2µg p75-Saporin (total dose) or PBS i.c.v. At 35 days a separate cohort of animals was injected (i.p) with either LPS (500µg/kg) or saline to confirm the cellular source of induced IL-1β. All procedures were performed, after ethical review by the Trinity College Animal Research Ethics Committee, under license from the Republic of Ireland National Regulatory Authority for animal experimentation (HPRA AE19136/P065) in accordance with Part 5 of the European Union (Protection of Animals Used for Scientific Purposes) Regulations 2012 (S.I. No. 543 of 2012). All biosensor work was performed under the same licence but with additional ethical approval from Maynooth University Research Ethics Committee (BSRESC-2017-002). All efforts were made to minimize the suffering and number of animals used.

### Immunohistochemistry

A subset of animals was transcardially perfused with heparinized saline and 10% formalin for choline acetyltransferase (ChAT) and IBA-1 immunohistochemistry. Coronal sections (10 µm) of paraffin wax-embedded tissue were cut on a microtome, at the level of the medial septum and hippocampus. Sections were labeled with goat anti-human ChAT (1/500; Millipore) or goat anti-IBA1 (Abcam), after pre-treatment with 1% H_2_O_2_/methanol (20 min), microwaving in citrate buffer (pH 6) for 2 × 5 min and pre-blocking with normal rabbit serum. IBA-1 sections were pretreated with 0.04% pepsin for 20 min before blocking. Thereafter, the ABC method was used as previously described (Cunningham *et al*., 2005).

To assess the extent of the cholinergic lesion, the sections were analysed and scored semi quantitatively in the medial septum. This ChAT score ranged from zero to four, with four being a normal medial septum with the expected number of medial septal ChAT-positive neurons and zero being a complete lesion with no visible medial septal Chat-positive neurons. In the control, PBS-injected mice, the medial septum was found to be normal, scoring a maximum of four. In the p75^NTR^-saporin mice, there was a substantial reduction in the number of medial septal neurons visible, however, there was variability in the extent of the lesion with scores ranging from zero to three (Suppl fig.1). IBA1 was quantified by calculation of the positive area on 8 bit images, converted to threshold binary images in order to readily quantify the IBA-1-positive area.

### Fluorescence-activated cell sorting (FACS) and flow cytometry (microglia, astrocytes)

#### Enzymatic digestion and myelin removal

Both hippocampi were carefully dissected and kept in 1 ml ice cold HBSS. The hippocampi were minced and dissociated in 5 ml of enzyme mixture containing collagenase (2 mg/ml), DNase I (28U/ml), 5% FBS, and 10 µM HEPES in HBSS, followed by a filtering step using a 70 µm cell strainer (BD Falcon) to achieve a single-cell suspension. Myelin from the single-cell suspension obtained was removed by subsequently incubating with Myelin Removal Beads II for 20min and passing through LS columns mounted over QuadroMACS magnet.

#### Cell sorting and Intracellular staining

The myelin-depleted single-cell suspension obtained above was incubated (15 min on ice) with anti-mouse CD16/CD32 antibody to block Fc receptors and subsequently incubated with anti-CD11b PEcy7 (Biolegend), anti-CD45 APC (Biolegend), and anti-GLAST PE (Milteny) antibodies for 30 min on ice. After a final wash with FACS buffer (1% BSA, 2 mM EDTA in PBS), cells were resuspended in 200 µl of FACS buffer and sorted on FACSAria Fusion cell sorter (Becton Dickinson) using 100 µm nozzle. 7AAD (Becton Dickinson) was used to gate out nonviable cells. Sorted cells were collected in 1.5 mL Lobind RNAse/DNAse free tubes containing 350 µL of sorting buffer (HBSS without Phenol Red supplemented with 7.5 mM HEPES and 0.6% glucose). Of note, 40–50,000 CD45lowCD11b + microglia and 80–100,000 GLAST+CD45-astrocytes were sorted (gating strategy described in Fig. 3A). Purity of sorted astrocytes and microglia was determined by quantitative polymerase chain reaction (qPCR) (Fig. 3B). For flow cytometric analysis of the cellular source of IL-1β, intracellular staining was performed on single cell suspensions according to instructions provided by True-Nuclear™ Transcription Factor Buffer Set (Biolegend). The antibodies used were anti-IL-1β PE (ThermoFisher), anti-GLAST APC (Milteny), anti-CD45 APCcy7 (Biolegend) anti-CD11b BV421(Biolegend) and anti-CD11c BV510 (Biolegend).

### Gene expression analysis

#### RNA extraction and cDNA synthesis from sorted cells and from homogenates

Sorted microglia and astrocytes samples were centrifuged at 10,000g for 10 min. Supernatant was carefully removed, and cell pellet was resuspended and lysed in 350 µl of RLT Plus Buffer containing β-mercaptoethanol. RNA extraction was done according to instructions provided by RNeasy Plus Micro Kit (Qiagen). cDNA was prepared from 10 to 50 ng RNA according to instructions of iScript™ cDNA Synthesis Kit (BioRad). Qiagen RNeasy^®^ Plus mini kits (Qiagen, Crawley, UK) were used for hippocampal and medial septum homogenates according to the manufacturer’s instructions. Samples were disrupted in 600 µl Buffer RLT using a motorized pestle followed by centrifugation at 14,800 rpm for 6 min through Qiagen Qiashredder columns to complete homogenization. The flow-through was collected and transferred to the genomic DNA (gDNA) Eliminator spin column and centrifuged at 14,800 rpm for 30 s. The column was discarded, and an equal volume of 70% ethanol was added to the flow-though and mixed until homogenous. Samples were placed in RNeasy mini spin columns in 2 mL collection tubes and centrifuged at 14,800 rpm for 15 s. On-column DNase digestion (Qiagen) RNase free DNase I incubation mix (80 µl) as an extra precaution to ensure complete removal of contaminating gDNA. RNA was well washed before elution with 30 µl of RNase-free water. RNA yields were determined by spectrophotometry at 260 and 280 nm using the NanoDrop ND-1000 UV–Vis Spectrophotometer (Thermo Fisher Scientific, Dublin, Ireland) and stored at −80°C until cDNA synthesis and PCR assay.

RNA was reversed transcribed to cDNA using a High Capacity cDNA Reverse Transcription Kit (Applied Biosystems, Warrington, UK). Four hundred nanograms of total RNA was reverse transcribed in a 20 µl reaction volume. Of note, 10 µl master mix (for each sample, master mix contained: 2 µl 10× RT Buffer; 0.8 µl 25× dNTP mix, 100 mM; 2 µl 10× RT random primers; 1 µl MultiScribe™ Reverse Transcriptase; 4.2 µl RNase-free water) was added to 10 µl RNA for each sample in a nuclease-free PCR tube (Greiner Bio-One, Monroe, NC). No reverse transcriptase and no RNA controls were also assessed by PCR. PCR tubes were placed in a DNA Engine^®^ Peltier Thermal Cycler PTC-200 (Bio-Rad Laboratories, Inc., Hercules, CA), and samples were incubated at 25°C for 10 min, 37°C for 120 min, and 85°C for 5 min (to inactivate reverse transcriptase). Samples were held at 4°C until collection and then stored at −20°C until assay.

#### Quantitative PCR

Reagents were supplied by Applied Biosystems (Taqman® Universal PCR Master Mix; SYBR® Green PCR Master Mix) and Roche (FastStart Universal Probe Master [Rox]; FastStart Universal SYBR Green Master [Rox]; Lewes, UK). For most assays, primers were designed using the published mRNA sequences for the genes of interest, applied to Primer Express™ software. Where possible, probes were designed to cross an intron such that they were cDNA specific. In some cases, the fluorescent DNA binding probe SYBR green was used in place of a specific probe. Primer and probe sequences, along with accession numbers for mRNA sequence of interest may be found in Table 1. Oligonucleotide primers were resuspended in 1× TE buffer (Tris Base 10 mM, EDTA 1 mM; pH 7.5–8.0) and diluted to 10 µM working aliquots. All primer pairs were checked for specificity by standard reverse transcription (RT)-PCR followed by gel electrophoresis, and each primer pair produced a discrete band of the expected amplicon size. For Taqman PCR, 24 µl of PCR master mix containing 12.5 µl Taqman^®^ Universal PCR Master Mix (or SYBR^®^ Green PCR Master Mix), 0.5 µl of each of the forward primer, reverse primer and probe, and 10 µL of RNase-free water was added to individual wells of a MicroAmp™ Optical 96-well Reaction Plate (Applied Biosystems, Warrington, UK). Where SYBR green was used, RNase-free water was substituted in place of the probe. To this, 1 µl of cDNA (equivalent to 20 ng of RNA) was added to each well to give a final reaction volume of 25 µL. Samples were run in the StepOne™ Real-Time PCR System (Applied Biosystems, Warrington, UK) under standard cycling conditions: 95°C for 10 min followed by 95°C for 10 s and 60°C for 30 s for 45 cycles. A standard curve was constructed from serial one in four dilutions of the cDNA synthesized from total RNA isolated from mouse brain tissue 24 hr after intra-cerebral challenge with 2.5µg LPS, known to upregulate target transcripts of interest in this study. A standard curve was plotted of Ct value versus the log of the concentration (assigned an arbitrary value since the absolute concentration of cytokine transcripts is not known). All PCR data were normalized to the expression of the housekeeping gene18s.

#### ELISA

Blood samples were centrifuged for 10 minutes at 1500*g* and plasma pipetted without disturbing the pellet. IL-1β ELISA kit (Biolegend) was used for determination in plasma and ELISA was performed according to manufacturers’ instructions. Briefly, 96-well plates were coated with 100 µl of capture antibody solution overnight at 4°C. The coated plate was washed with 300 µl wash buffer per well four times and then tapped upside down onto absorbent paper to ensure complete removal of contents. The plates were blocked by adding 200 µl assay diluent before sealing and incubating for one hour at room temperature with shaking. After four washes, 100µl diluted samples and standards were added to their respective wells and incubated for two hours at room temperature with shaking. After four washes, 100µl of detection antibody solution was added to each well and incubated for one hour at room temperature with shaking. The plate was then washed four times and 100µl Avidin-HRP solution was added to each well and incubated for 30 minutes at room temperature with shaking. The plate was then washed five times, 100 µl TMB Substrate Solution was added to each well on the plate and then incubated in the dark for 15-30 minutes or until appropriate colour development across the concentration range of cytokine standards. 100µl Stop Solution (H_2_SO_4_) was added to each well on the plate. For brain ELISA, IL-1β Quantikine kits (R&D Systems) were used. Hippocampal samples were weighed and homogenised in homogenisation buffer using a motorised pestle (150 mM NaCl, 25 mM Tris-HCl, 1% Triton X100, pH 7.4). A DC (detergent compatible) protein assay was performed prior to ELISA. 200 µl reagent B was added to each well and the plate gently shaken to mix. Plate was read at 750 nm after 15 minutes. ELISA standard, substrate solution, wash buffer and control were prepared as per manufacturer’s instructions. 50 µl assay diluent was added to each well of the microplate. 50 µl sample or standard was added to each well before the plate was sealed and incubated at room temperature for 2 hours. The plate was washed as previously described with wash buffer (400 µl) 5 times. 100µl antibody conjugate was added to each well before the plate was sealed and incubated at room temperature for 2 hours. The plate was again washed as above, 5 times. 100µl substrate solution was added and incubated for 30 minutes in the dark. 100 µl stop solution was added and the plate tapped gently to mix. The absorbance reads of both plasma and hippocampal samples were taken at 450 nm and 570 nm within 15 minutes on a BioTek Synergy HT Multi-Detection Microplate Reader.

#### Choline biosensor preparation

Changes in extracellular choline were monitored in freely-moving mice using constant potential amperometry (CPA) with choline microelectrochemical biosensors (Baker *et al*., 2015; Teles-Grilo Ruivo *et al*., 2017; Baker *et al*., 2018). In brief, biosensors were constructed from Teflon^®^-coated, Pt/Ir (90%/10%) wire (75µm bare diameter, 112 µm coated diameter; Advent Research Materials). One end was stripped of Teflon^®^ insulation and soldered into a gold-plated pogo pin (Bilaney Consultants Ltd.). A fresh disk was cut at the opposite end, which acted as the active surface. The disk surface was coated with a layer of electropolymerized poly-*o*-phenylenediamine (PPD; >98%), a well characterised interference rejection layer making the sensor highly selective for choline (Lowry *et al*., 1998; Baker *et al*., 2015; Baker *et al*., 2018). The PPD-modified electrode was initially dipped in methylmethacrylate (MMA; 99%) and cellulose acetate solutions, and then sequentially dipped into choline oxidase (ChOx; from Alcaligenes sp., EC 232-840-0), bovine serum albumin (BSA; fraction V from bovine plasma), glutaraldehyde (Grade 1, 25%) and polyethyleneimine (PEI; 80% ethoxylated) using a dip absorption method. This process was repeated a total of 10 times, allowing a 4-minute drying period between layers, producing a platinum/PPD-polymer composite (PC)/ChOx-modified electrode (Pt/PPD-PC/ChOx/PC). The sensors were dried for a minimum of 1 hour at room temperature and stored at 4°C before use.

Prior to implantation, all biosensors were calibrated *in vitro* in a standard three-electrode glass electrochemical cell in 20 ml PBS (pH 7.4). A saturated calomel electrode (SCE) acted as the reference electrode and a bare Pt wire served as the auxiliary electrode. CPA (+700 mV (choline)) was performed in all electrochemical calibration experiments, using custom designed, low-noise potentiostats (Biostat IV, ACM Instruments) with a notebook PC, a PowerLab AD (AD Instruments Ltd.) interface system and LabChart^®^ for Windows (v8, AD Instruments Ltd.) or an eDAQ e-corder and eDAQ Chart v5.5.23 (Green Leaf Scientific). Biosensors were allowed to settle under the influence of the applied potential until the non-faradaic current reached a stable baseline. To validate, choline concentration was sequentially increased from 0 to 3 mM by adding aliquots of choline chloride (ChCl), followed by a brief (ca. 20s) stirring after each aliquot. The lower limit of detection of the choline biosensors was 100 nM (Baker *et al*., 2018). Biosensors were selected for implantation if the calibration current values were not significantly different to the average.

#### Surgical implantation of choline biosensors

Mice were anaesthetised using the volatile anaesthetic, isoflurane (4% at 450 ml/min in air for induction, 0.9-2.5% at 250 ml/min in air for maintenance, Isoflurin^®^) using a Univentor 400 Anaesthetic Unit (Agnthos). Once surgical anaesthesia was established, the upper head was shaved, mice were positioned in a stereotaxic frame, administered a s.c. injection of Buprecare (0.05 mg/kg), a s.c. of lidocaine along upper surface of the head and connected to a pulse oximeter (MouseSTAT® Jr., Kent Scientific). Under sterile conditions, the skull was exposed and cleared of overlying fascia. The head was levelled between bregma and lambda, and craniotomies were drilled using a 0.7 mm steel burr (Fine Science Tools) for i.c.v. injections (see *surgery and animal treatments*) and biosensor implantation in the following coordinates; dorsal hippocampus (AP −2.2, ML +1.8 mm, DV −1.75 mm) and medial prefrontal cortex (AP +1.95 mm, ML −0.30 mm, DV −1.90 mm) (AP/ML coordinates with respect to bregma, DV coordinates from the surface of the brain). Additional craniotomies were drilled for a reference electrode and three support screws (BASi^®^), one of which was wrapped with the auxiliary electrode. Under stereotaxic guidance, the choline biosensors and reference electrode were slowly implanted and fixed in place using dental acrylate (Dentalon^®^, Heraeus-Kulzer). The electrode pogo pins were inserted into a Delrin^®^ 12-channel pedestal (Bilaney Consultants Ltd.), which was subsequently secured to the skull using dental acrylate. Post-surgery, mice were administered a s.c. injection of sterile saline and allowed to recover in a thermostatically controlled cage (Datesand). Animals were assessed daily for good health and allowed a minimum of 3 days before being connected to instrumentation for *in vivo* recording.

#### In vivo biosensor recording

Mice were singly housed in motion controlled Raturn^®^ sampling cage systems (BASi). The head-mounted Delrin^®^ pedestal was connected to the potentiostat (Electrochemical and Medical Systems, EMS) using a lightweight, flexible six-core cable. This set-up allowed free movement of the animals during biosensor recording. Following the application of an applied potential (+700 mV), mice were allowed a further 20+ hours to ensure background biosensor current has settled to a stable baseline. All amperometric recordings from each working electrode (biosensor) channel was recorded at 1 kHz, and a PowerLab interface system was used for analog/digital conversion before the data was collected on a Mac running LabChart.

#### Statistical Analysis

Data were assessed for normality and thereafter analyses of p75-saporin versus vehicle were performed by Student’s t-test. All subsequent experiments with a 2×2 design were analysed by 2-way ANOVA using lesion group and acute treatment as factors. Subsequent experiments with multiple treatments were assessed by 1-way ANOVA using treatment as the only factor. All data-sets were analysed using GraphPad Prism v9.0.

## Results

### Loss of medial septal cholinergic neurons triggers hippocampal microglial activation

Intracerebroventricular injection of p75-saporin (0.6µg bilaterally, i.c.v) produced a partial lesion of medial septum cholinergic neurons, as is evident by ChAT immunostaining (Fig. 1A&B). This also resulted in significant loss of ChAT-positive synaptic terminals in the hippocampus (CA1) (Fig. 1C). Robust microglial activation was evident in the hippocampus using IBA-1 immunostaining (Fig. 1D-E). Quantification of IBA-1 positive area in hippocampi of p75-saporin-injected mice showed increased IBA-1-positive area compared to PBS-injected group (Fig. 1F; p<0.0001). There was an inverse correlation (R^2^=0.47, p<0.0001) between the medial septum ChAT score and hippocampal IBA-1-positive area (Fig. 1G). This increased microglial activation was concomitant with the increase in microglial number, as evident by IBA-1 immunostaining (Fig. 1D) and increased CD45^low^CD11b^+^ microglia (Fig. H&I, p<0.0001) in hippocampi of lesioned animals. A time course across 35 days, in the medial septum and hippocampi of lesioned animals and controls, showed strong induction of pro-inflammatory genes in the days after p75-saporin injection, with significant inter-individual variation. Inflammatory cytokines in the hippocampi showed an early and prominent increase in mRNA expression of *Tnfa* at both 3 (p<0.05) and 7 days (p<0.05) post lesion and it’s levels remained somewhat elevated (though not significantly) even at 35 days post-lesion. After an early increase, *Il1b*, in hippocampi returned to control levels by 14 days post lesion. *Sra2*, a scavenger receptor gene, also showed marked increase at 3 days (p<0.001) and though it decreased by 7 days it still remained elevated with respect to controls for 35 days. The scavenger receptor *Cd68* and *Trem2* (Triggering Receptor Expressed on Myeloid Cells 2; a determinant of microglial phenotype and clearance of debris (Hickman & El Khoury, 2014; Keren-Shaul *et al*., 2017) both showed slower and modest increases but both remained elevated at 35 days (Fig. 1J). Similar to what was observed in the hippocampus, levels of *Tnfa* (p<0.05) *and Il-1b* (p<0.001, p<0.05) were elevated in medial septum until 7 days post-lesion. However, in general, microglial markers (*Sra2, Cd68, Trem2*) returned to basal levels by 14 days and only *Tnfa* remained significantly elevated at 35 days (Fig. 1K). Therefore, those fields that have been depleted of cholinergic tone show more persistent evidence of microglial activation than the region of the neurons from which that ACh originates.

**Figure 1:**
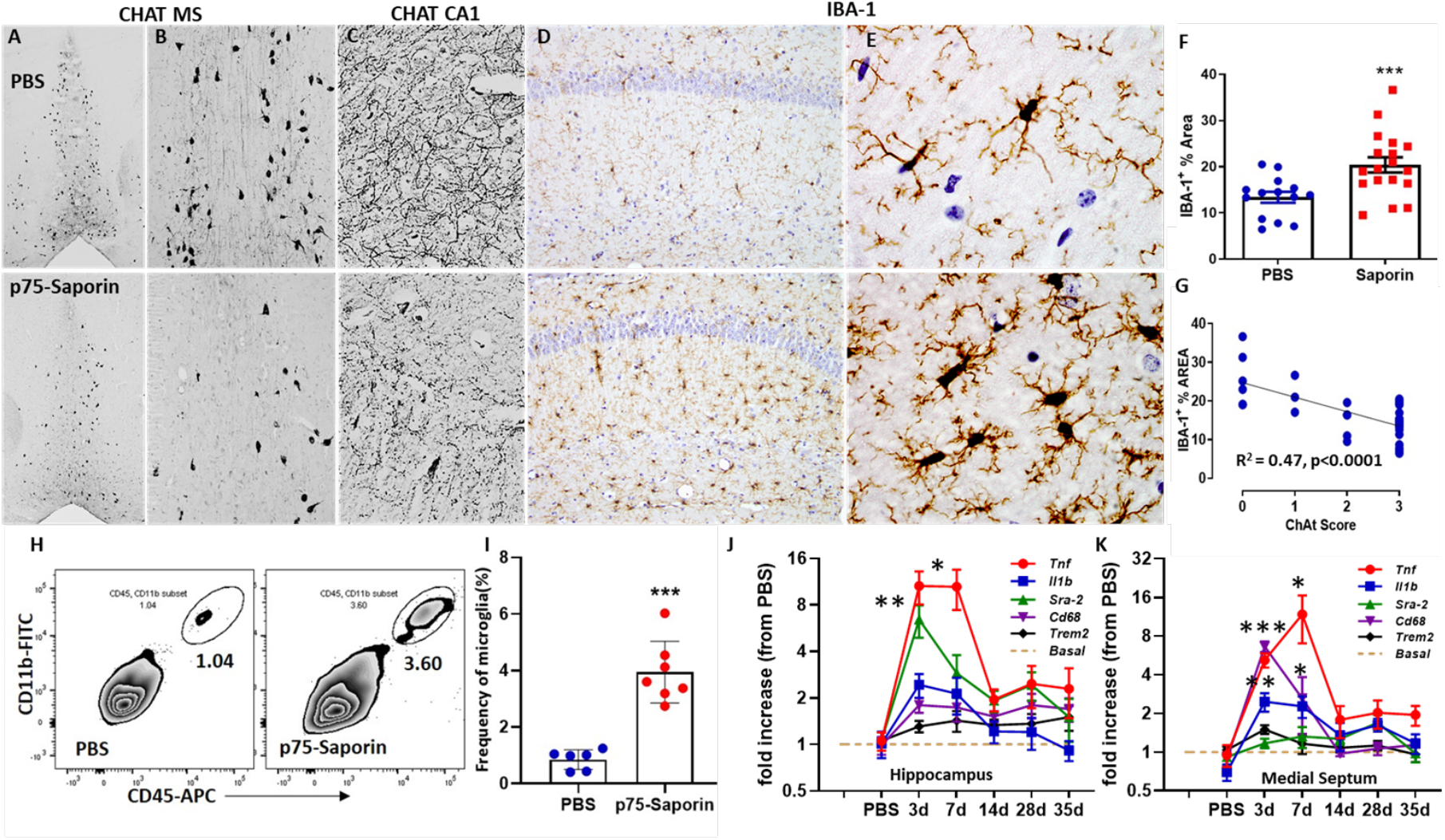
Microglial activation and temporal changes in proinflammatory markers in hippocampus and medial septum post cholinergic lesion. A,B) Representative photomicrographs of ChAT-positive neuronal depletion in the medial septum/diagonal band in 1.2 µg p75-Saporin-injected mice, scale bar: 200 µm. C) Photomicrograph showing ChAT-positive terminals at the level of hippocampus of the same animal, scale bar: 500 µm. D,E) Image showing IBA-1 labelling in hippocampal CA1 region of PBS and p75-Saporin injected animals at 20X and 100X. F) Graph showing percent IBA-1+ area in hippocampi of PBS and p75-Saporin injected animals (n=20 PBS, n=26 saporin, p=0.0002, t-test). G) Graph depicting inverse correlation between IBA-1+ area (%) and medial septum ChAT score (R^2^=0.47, p<0.0001, linear regression analysis). H,I) Representative flow cytometry graphs showing frequency of CD45lowCD11b+ microglia in PBS and p75-saporin injected animals (n=6 PBS, n=7 saporin, p=0.001, t-test). J,K) Time courses of hippocampal and medial septum mRNA expression by qPCR of p75-saporin (1.2µg) lesioned mice and PBS controls. *Tnfa, Il1b, Sra2, Cd68* and *Trem2*, n = 5 for each group except at 14 days where n = 4. One-way ANOVA with Bonferroni post-hoc test at 3, 7,28 and 35 days (* = P<0.05, ** = P<0.01,*** = P<0.001). All data are represented as mean ± SEM (F-K).

### The hypocholinergic hippocampus produces exaggerated inflammatory responses to systemic LPS

We next investigated whether the hypocholinergic hippocampus was hypersensitive to subsequent systemic inflammatory challenges. In an earlier study we found that p75-saporin, at 0.08 µg i.c.v, did not lead to priming of either hippocampal or medial septal microglia (Field *et al*., 2012). However, those experiments were performed with a deliberately low dose of the toxin, producing a limited lesion of <30% and tissues were examined 40 days post-lesion, making it possible that primed microglia had reverted to their basal phenotype or that limited cholinergic loss was insufficient to prime microglia in the first instance. Thus we investigated (Fig. 2A) whether microglia were primed by (1) a more severe lesion of the basal forebrain cholinergic system (1.2 µg p75-Saporin, 40 days post-lesion, henceforth called **high dose, 40d**) or (2) by examination of the inflammatory response to a lower dose, but at an earlier time-point following lesion (0.4 µg p75-Saporin, 14 days post-lesion, henceforth called **low dose, 14d**). IL-1β protein expression was induced by systemic LPS in the hippocampus of p75-Saporin-lesioned animals to a significantly greater degree than in non-lesioned, LPS-treated animals; both with low dose, 14d (Fig. 2B) and with high dose, 40d (Fig. 2D). 2-way ANOVA showed an interaction of cholinergic lesion and LPS for both low dose (*F=4*.*698, df*_*1,13*_, *p=0*.*0494*) and high dose lesions (*F*=7.83, df_1,14_, *p*=0.0142) and p75-saporin+LPS was significantly different to PBS+LPS (Bonferroni post-hoc, p<0.05, <0.001 for low and high dose respectively).

**Figure 2:**
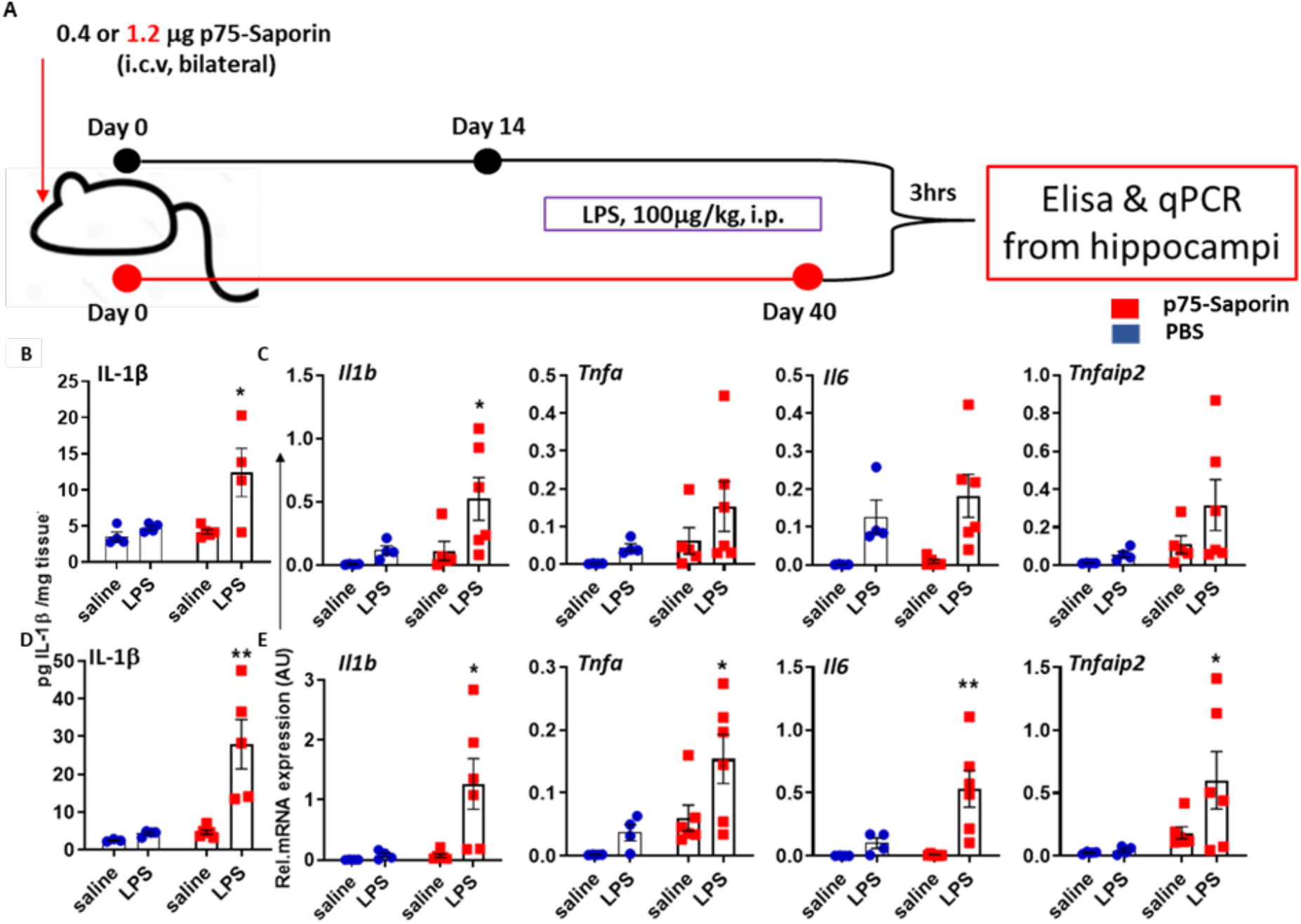
LPS-induced inflammation in the hippocampus of p75-saporin-lesioned animals. A) Schematic showing tissue sampling 3hours post LPS (100 µg/kg) treatment in 14 days after 0.4 µg p75-sap mediated lesion (black) and 35 days after 1.2 µg p75-sap mediated lesion (red). B) Exaggerated hippocampal IL-1β protein levels determined by ELISA 3 hours after LPS treatment in 14 days post lesioned mice (0.4 µg). C) Graphs representing changes in the expression of *Il1b, Tnfa, Il6* and *Tnfaip2* mRNAs 3 hours after LPS treatment in 14 days post lesioned (0.4 µg) mice. D) Graph showing hippocampal IL-1β protein levels determined by ELISA 3 hours after LPS treatment in 40 days post lesioned mice. E) Graphs representing changes in the expression of hippocampal *Il1b, Tnfa, Il6* and *Tnfaip2* mRNAs 3 hours after LPS treatment in 40 days post lesioned (1.2 µg) mice. All data are plotted as mean ± SEM with n = 4 (Saline in PBS), n = 4 (LPS in PBS), n = 6 (Saline in p75-saporin) and n = 6 (LPS in p75-Saporin) and analysed by two-way ANOVA with Lesion and treatment as between subjects factors. Significant differences are denoted *p < 0.05, **p < 0.01, and ***p < 0.001 by Bonferroni post-hoc analysis.

**Figure 3:**
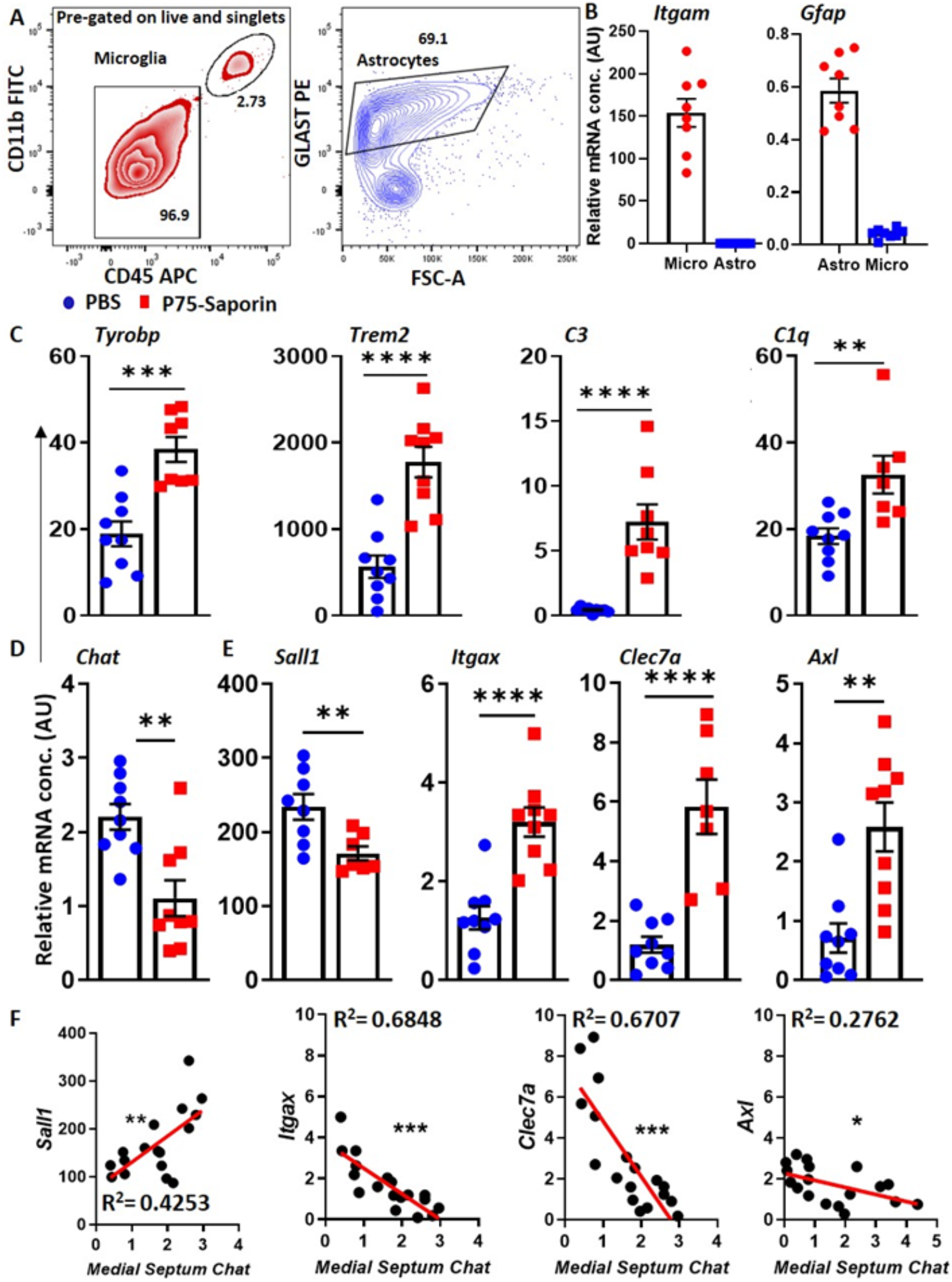
Characterisation of FACsorted microglia from lesioned and normal animals. A) Graphs depicting gating strategy used for FACsorting microglia and astrocytes. B) Purity of isolated cells was determined by qPCR for *Gfap* and *Itgam* in both astrocytes and microglia together. C) Graphs showing the changes in the expressions of *Tyrobp, Trem2, C3* and *C1q* in sorted microglia from the hippocampi of lesioned (1.2 µg p75-Saporin) and PBS injected mice (C) (** p=0.0058, *** p=0.0002, **** p<0.0001, t-test). D) Graph showing medial septum *Chat* in lesioned (1.2 µg p75-Saporin) and PBS-injected mice. E) Graphs showing expressions of *Sall1* (marker of resting state), *Itgax, Clec7a* and *Axl* (primed microglial signature) in isolated microglia from lesioned and PBS-injected mice 35 days after injection (**p=0.0058, *** p=0.0002, **** p<0.0001, t-test, Data are presented as the mean ± SEM). F) Regression analysis of *sall1, Itgax, Clec7a*, Axl expression vs.medial septum *Chat* mRNA expression in the same animals (*p < 0.05, **p < 0.01, and ***p < 0.001, after linear regression).

The exaggerated IL-1β effect was also apparent at the mRNA level. Two-way ANOVA analysis of *Il1b* revealed a main effect of cholinergic lesion (*F=4*.*84, df*_*1,15*_, *p=0*.*0439*), and of LPS (*F*=8.59, *df*_*1,15*_ *p*=0.0103) in low dose lesioned animals and similar effect of lesion (*F*=5.58, *df*_*1,16*_, *p*=0.0312) and of LPS (*F*=4.64, *df*_*1,16*_ *p*= 0.048) in high dose lesioned animals. Bonferroni post-hoc revealed significant differences between lesion+LPS vs PBS+LPS at both doses, *p*<0.05 (Fig. 2C&E). Effects on the other pro-inflammatory genes examined showed broadly similar patterns but were less robust. Systemic LPS effects on *Tnfa, Il6 and Tnfaip2* were not significantly different in the hippocampus of low dose p75-Saporin lesioned animals compared to vehicle lesion controls (Fig. 2C). However, systemic LPS did induce exaggerated expression of *Tnfa* (*p*<0.05), *Il6* (*p*<0.01) and *Tnfaip2* (*p*<0.05) in the hippocampus of p75-saporin animals compared to LPS in non-lesioned animals (Fig. 2E).

### Cholinergic depletion shifts microglia from homeostatic to primed state

In figure 2 we show exaggerated IL-1 responses in the hippocampus and suggest that the microglia may be primed. Therefore we sorted microglial cells by FACS in order to further examine their phenotype in the absence of LPS. After sorting by FACS using CD11b/CD45 for microglia and GLAST1 for astrocytes (Fig. 3A), we verified purity of these populations using PCR for *Itgam* and *Gfap* respectively. Microglia preparations showed undetectable levels of Gfap transcripts (Figure 3B). Microglia from lesioned animals showed elevated expression of *C3, C1q, Trem2, Tyrobp* (Fgure 3C, p<0.0001, <0.01, <0.0001, <0.001 respectively). These data indicate that microglia have diverged from their homeostatic state.

*Sall1* has been identified as a key transcriptional regulator of microglial identity and to be important for maintenance of its homeostatic state. Ablation of *Sall1 results* in conversion of microglia into inflammatory phagocytes (Buttgereit *et al*., 2016). Moreover, weighted Gene Co-expression Network Analysis (WGCNA) of microglia gene expression signature by aging and neurodegenerative conditions (Holtman *et al*., 2015) has revealed the gene expression signature of microglial priming, showing a number of key signature genes shared across multiple models. We therefore analysed the expression of *Sall1* and the priming ‘hub’ genes *Clec7a, Itgax* and *Axl*, in microglia sorted from hippocampi of PBS and p75-Saporin (1.2 µg p75-Saporin, 35 days post-lesion) animals. We found the expression of *Sall1* (p=0.0097) to be significantly reduced in microglia isolated from the hippocampi of p75-Saporin treated animals compared to PBS controls (Fig. 3E). Moreover, we found a significant positive correlation between the expression of hippocampal *Sall1* and medial septal *Chat* expression (R^2^=0.4253, p<0.001; Fig 3F). Consistent with loss of septal Chat predicting loss of microglia homeostasis, we found significant upregulation of *Clec7a* (p<0.0001), *Itgax* (p<0.0001) *and Axl* (p<0.0012) in microglia isolated from hippocampi of PBS and p75-Saporin animals (Fig. 3E) and the expression of *Clec7a* (R^2^=0.6707, p<0.0001) and *Itgax* (R^2^=0.6848, p<0.0001) were strongly inversely correlated with their corresponding medial septal *Chat* expression. *Axl* showed weaker, but significant inverse correlations to septal *Chat* expression (R^2^=0.2762, p<0.05) (Figure 3F).

### Microglia, not astrocytes, produce significant IL-1β after cholinergic depletion

To demonstrate the cellular source of IL-1β post-systemic inflammation with LPS, we performed intracellular staining and subsequent flow cytometry on single cell suspensions obtained from hippocampi of lesioned and normal animals, challenged with LPS or saline. The data show significant (p=0.0095) IL-1β labelling in the microglia from the lesioned group that received LPS compared to controls (Fig 4A-G). IL-1β production was limited to microglia and astrocytes did not appear to contribute to this expression (Fig. 4 J,K). Moreover, the microglial priming marker CD11c was more highly expressed in lesioned animals (Fig 4H) and its expression was directly correlated to the expression of IL-1β among LPS-treated animals (Fig. 4I; Pearson’s correlation analysis: R^2^=0.7298, p<0.0001).

**Figure 4:**
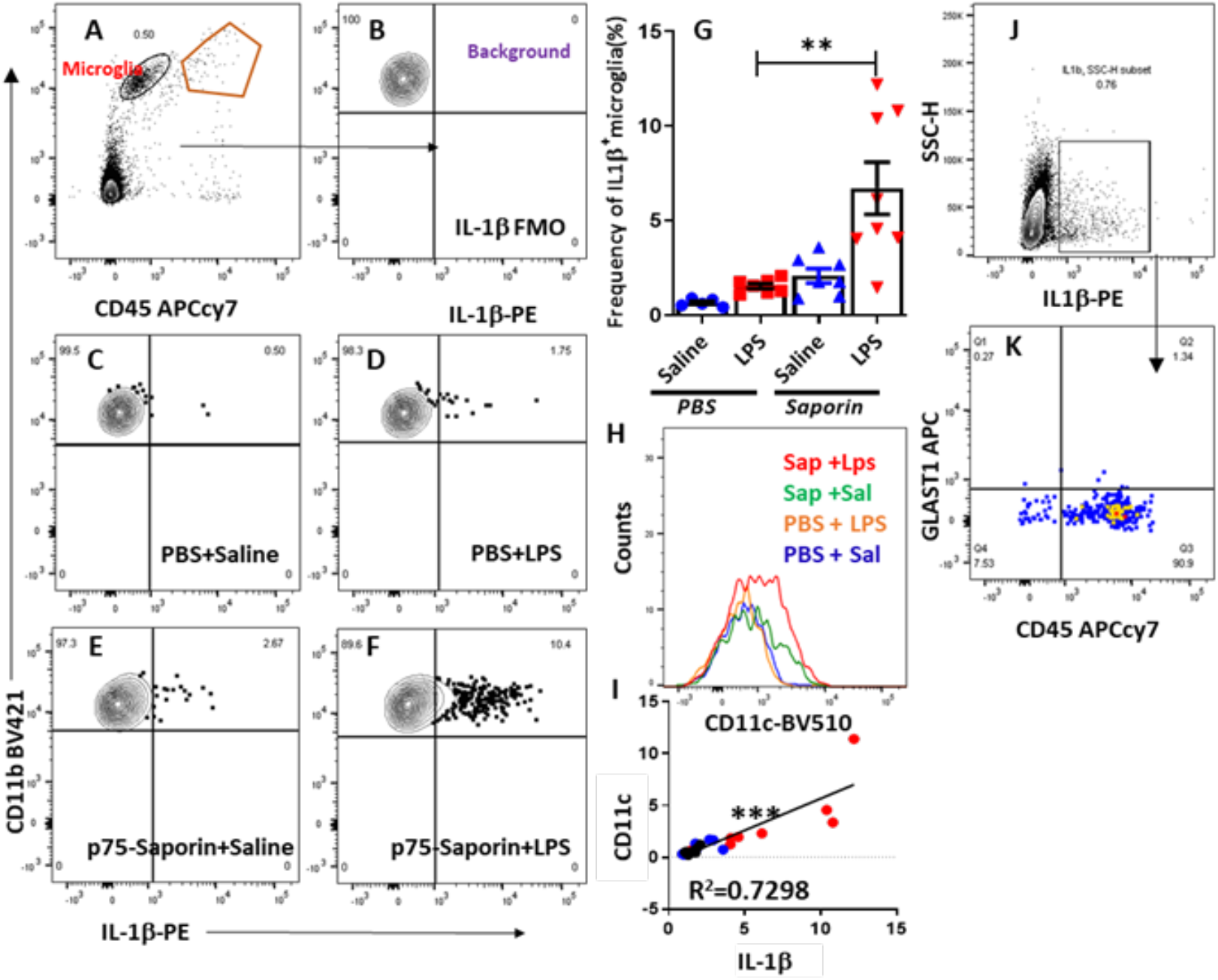
Microglia, not astrocytes, are the producer of IL-1β after LPS treatment of lesioned animals. Flow cytometry dot plot showing microglial gating strategy (A). Graphs depicting IL-1β production from gated microglial cells in all the treatment groups (LPS: 500 µg/kg), FMO control for PE channel contains all antibodies except anti IL-1β PE (B-F). Graph showing frequency (%) of IL-1β producing microglial cells in all the treatment groups (G). Histogram depicting expression of CD11c on gated microglia belonging to all the treatment groups (H). Strong correlation (R^2^=0.7298) between IL-1β producing microglia and the level of expression of CD11c on their surfaces (I) Graph showing total IL-1β producing cells are negative for GLAST-1 (astrocyte marker) and positive for CD45 (J-K) (**p=0.0058, ***p=0.0002, **** p<0.0001, t-test, Data are presented as the mean ± SEM).

### Microglia express the α7-nAChR, and nicotinic agonism suppresses microglial Il1b

As mentioned above, microglia have been shown to express α7-nAChR and to respond to nicotinic agonism but the most convincing evidence for this is *in vitro* (Shytle *et al*., 2004; De Simone *et al*., 2005). In order to access the expression of α_7_-nAChR on microglia and astrocytes from normal mice in *in vivo* experiments, we first sorted microglia (CD45^low^CD11b^+^) and astrocytes (GLAST-1^+^) from the hippocampal single cell suspension. Sorted microglia and astrocytes were then stained with anti-α_7_-nAChR antibody and analyzed by flow cytometry. We found approximately 70% of CD45^low^CD11b^+^ microglia were positive for α_7_-nAChR. Astrocytes also showed some expression of α_7_-nAChR, though significantly fewer (1%) when compared to microglia (Fig. 5A-B).

**Figure 5:**
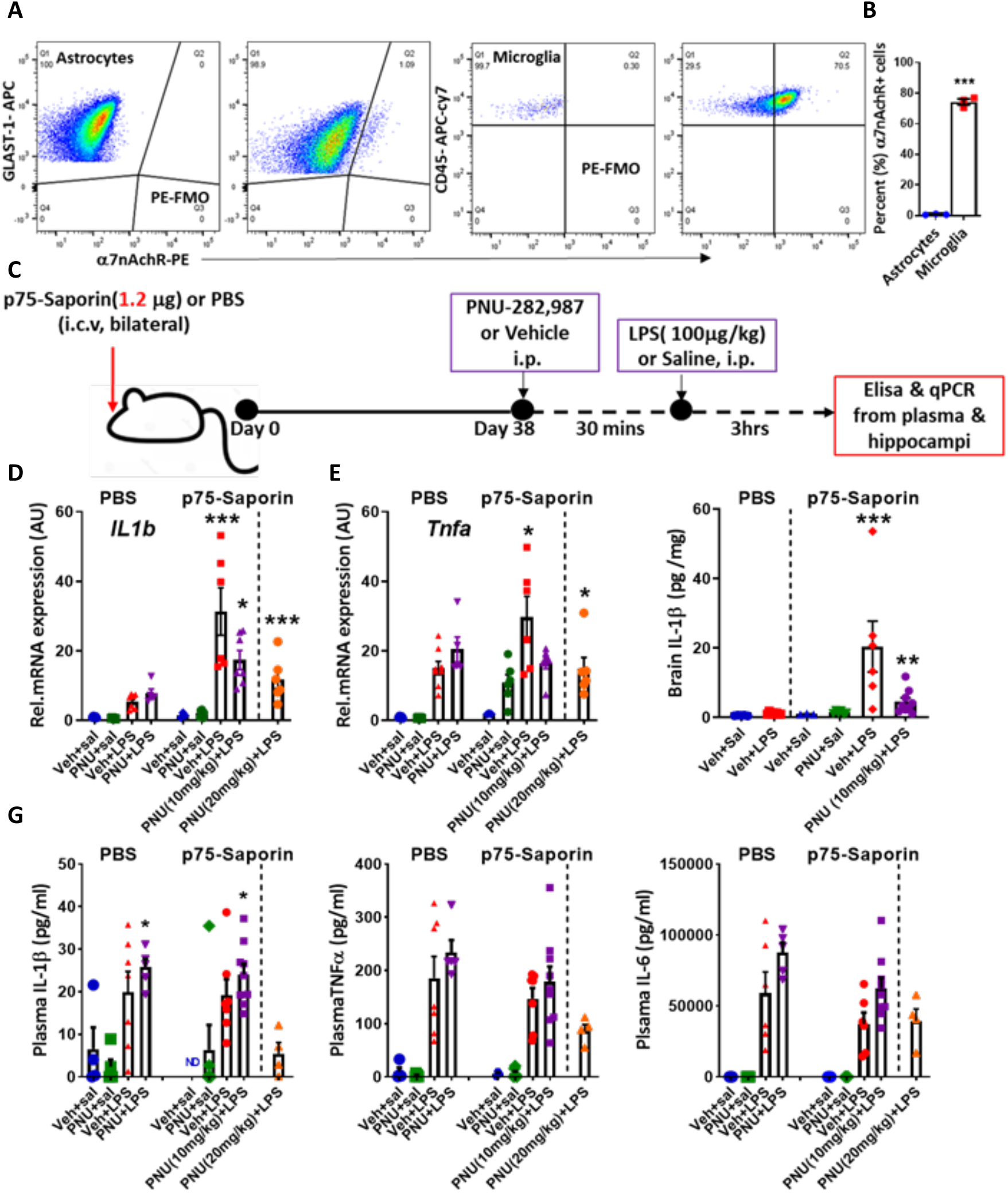
Microglial expression of and regulation by α7-nAChR cholinergic receptors. Flow cytometry dot plots showing α7-nAChR expression on FACsorted microglia and astrocytes with their corresponding FMO controls (A). Graph showing frequency (%) of α7-nAChR+ microglia and astrocytes (***p < 0.001, t-test, mean ± SEM) (B) Schematic showing timeline of PNU-282,987 administration, LPS injection and tissue sampling in lesioned (1.2µg) and PBS control mice (C). Hippocampal expression of *Il1b* and *Tnfa* by qPCR of lesioned mice versus PBS controls at 38 days [Mice administered LPS (100µg/kg) or saline and PNU-282,987 (PNU) at one of two doses (10mg/kg or 20mg/kg) or vehicle control (veh)] (D-E). Hippocampal brain IL-1β protein expression by ELISA of lesioned mice versus PBS controls at 38 days, administered LPS or saline (sal) and PNU or veh (F). Plasma IL-1β, TNF-α and IL-6 levels determined by ELISA of lesioned mice versus PBS controls at 38 days, administered LPS or saline (sal) and PNU or veh(G). Analysed by ordinary One-way ANOVA, Significant differences are denoted *p < 0.05, **p < 0.01, and ***p < 0.001 by Bonferroni post-hoc analysis.

Since microglia bear the α_7_-nAChR receptor and become primed upon cholinergic denervation, we hypothesised that exaggerated responses to systemic LPS would be suppressed by nicotinic cholinergic agonists and that microglial cells may be particularly responsive to these treatments since they had been depleted of cholinergic enervation. We used the brain penetrant nicotinic agonist PNU282,987 (Walker *et al*., 2006) and sought a dose that would not block systemic responses to low dose LPS. Importantly *c*irculating levels of IL-1β, TNF-α and IL-6 levels were not increased by prior cholinergic lesion (p75-saporin+veh; Figure 5 G). These cytokines were induced by systemic LPS: IL-1β (not significant), TNF-α (p=0.0023) and IL-6 (p=0.0011) compared to saline-treated, but LPS-induced levels were not altered by prior cholinergic lesion (p75-saporin+LPS vs veh+LPS, Figure 5 G). Moreover, the nicotinic agonist PNU282,987 (at 10 mg/kg) did not affect levels of these secreted cytokines. Given the previously described suppression of LPS-induced inflammation, albeit at the much higher LPS dose of 6 mg/kg (Borovikova *et al*., 2000; Wang *et al*., 2003), it was important to show that at 20 mg/kg PNU282.987 did suppress LPS-induced circulating cytokines, although not statistically significantly (Fig. 5 G).

The effects of LPS and PNU-282,987 on pro-inflammatory cytokine mRNA expression were investigated in the hippocampi of p75-saporin lesioned mice and PBS controls at 38 days post-lesion.

LPS significantly increased the expression of *Il1b* mRNA in p75-saporin treated animals compared to PBS controls (p<0.001, Bonferroni post-hoc, 1-way ANOVA) confirming the priming effect of p75-saporin lesion. Both 10 and 20 mg/kg doses of PNU were effective in reducing the expression of *Il1b* mRNA due to LPS in p75-saporin treated animals (p<0.05, p<0.001, Bonferroni post-hoc, 1-way ANOVA) (Fig. 5D). Similar significant increase in *Tnfa* mRNA (p<0.05, Bonferroni post-hoc after significant One-way ANOVA) was observed in LPS-injected lesioned animals compared to PBS controls and both doses of PNU-282,987 were effective in reducing LPS-induced transcription of *Tnfa* in p75-saporin treated animals (p<0.05, Bonferroni post hoc, 1-way ANOVA) (Fig. 5E). Hippocampal IL-1β protein was determined by Quantikine ELISA assay and was significantly elevated in p75-saporin-lesioned LPS-treated animals, compared to PBS controls (p<0.001, Bonferroni post-hoc, One-way ANOVA). PNU-282,987 significantly reduced IL-1β levels by LPS in p75-saporin animals (P<0.05, Bonferroni post-hoc, One-way ANOVA) (Fig. 5F). Importantly this nicotinic agonist had no impact on the transcription of these genes by LPS in non-lesioned animals. These data indicate that the withdrawal of basal forebrain cholinergic tone leaves the hippocampus hypersensitive to subsequent inflammatory stimulation and that the application of an α_7_-nAChR agonist is anti-inflammatory specifically in that context.

The observed suppression, by PNU-282,987, of the exaggerated cytokine response in p75-saporin+LPS mice (Figure 5) may simply reflect the downregulation of the acute LPS-induced cytokine response or may arise through effects of nicotinic agonism on the underlying microglial state, altering microglial priming and thus affecting their propensity to show the exaggerated responses to systemic LPS. To address this question we performed 2 different experiments: 1) We assessed whether nicotinic agonism may suppress acute inflammation by interrogating whether LPS may produce effects on acetylcholine dynamics in the hippocampus and whether those dynamics were altered in lesioned animals; 2) We assessed whether nicotinic agonism was sufficient to alter the microglial primed state even in the absence of LPS, in order to assess whether PNU282,987 effects were or were not contingent on direct effects on LPS signalling.

### LPS induces dynamic changes in hippocampal and cortical acetylcholine

First, we hypothesised that LPS may directly influence hippocampal ACh levels. We used custom-designed choline biosensors to monitor dynamic changes in choline, as a proxy for ACh, in the hippocampus, in real-time before and during LPS treatment. Continuous choline biosensor current (*I*_Choline_) recordings in PBS control and p75-saporin (1.2µg) lesioned mice following the systemic administration of LPS (500µg/kg) or sterile saline (0.9%) showed that LPS produced a statistically significant increase in the area under the curve for choline current (Figure 6). Data were analysed by two-way ANOVA and showed a significant interaction of lesion and treatment and Bonferroni post-hoc tests showed significant differences between LPS and saline only in non-lesioned animals (denoted *p=0.0314; **p < 0.01). This increase occurred within minutes of LPS challenge, in both hippocampus (6A) and frontal cortex (6D), and persisted across several hours. This LPS-induced increase was almost completely ablated in p75-saporin lesioned animals, in both regions (Figure 6 C,D). Therefore, LPS acutely increases hippocampal and frontal cortical ACh levels, which originates in the basal forebrain and is absent in lesioned animals. This persists across the period of acute inflammatory activation in these regions.

**Figure 6:**
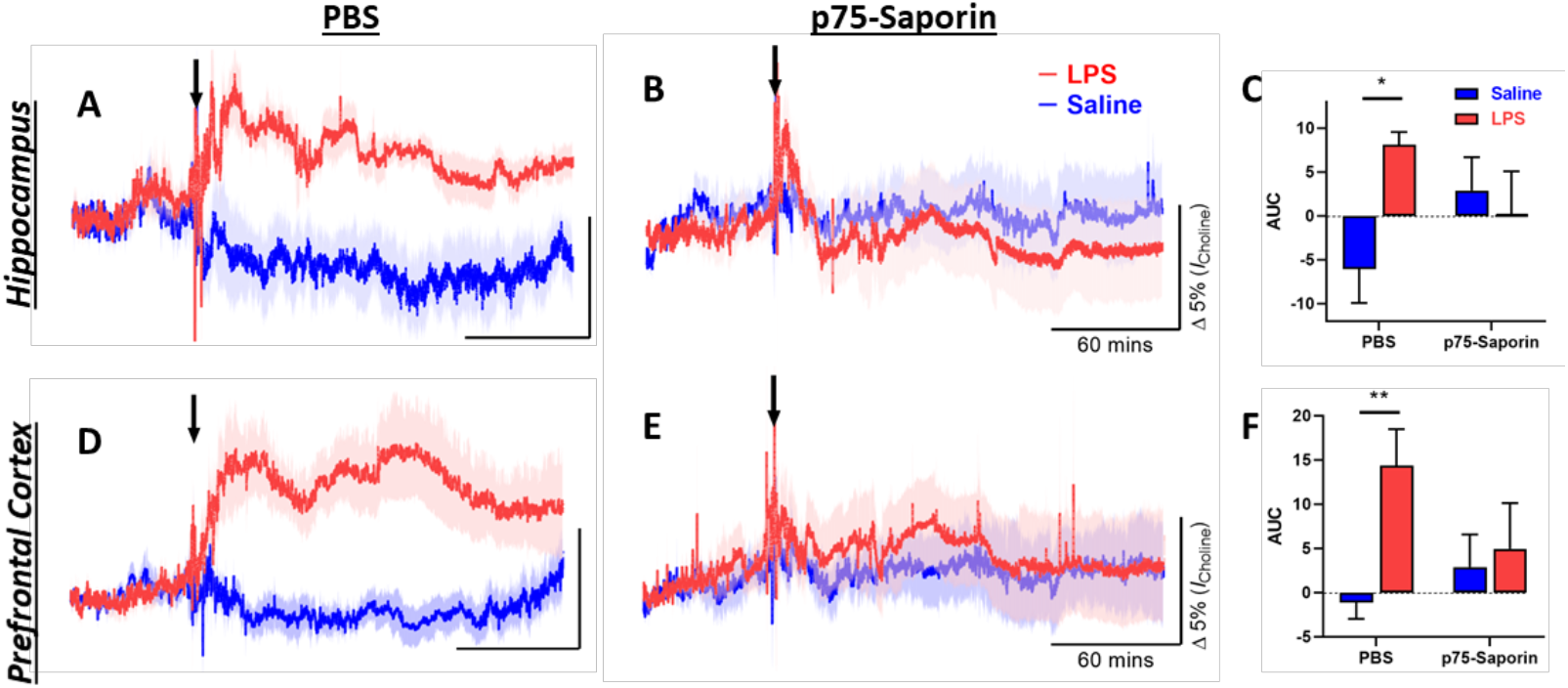
LPS-induced changes in extracellular choline. Continuous choline biosensor current (*I*_Choline_) recordings in PBS control and p75-saporin (1.2µg) lesioned mice following systemic administration (i.p.) of LPS (500µg/kg; red) or saline (0.9%; blue). Temporal profile of hippocampal (A-B) and prefrontal cortex (D-E): *I*_Choline_ changes in both regions. Data are expressed as mean (±SEM), normalised to a 1-hour pre-injection baseline and presented as relative percentage change in recorded *I*_Choline_. Area under the curve (AUC) analysis of *I*_Choline_ Data analysed by 2-way ANOVA, significant differences by Bonferroni post-hoc analysis are denoted *p=0.0314; **p=0.0048. C, hippocampus: n=6 for PBS+saline n=7 for all others. F, frontal cortex: PBS-saline (n=10) versus PBS-LPS (n=10).

### Multiple PNU-282,987 treatments attenuate priming of microglia by cholinergic lesion

Second, lesioned mice from day 25 onwards (giving adequate time for microglia to be primed) were treated once daily with PNU (10mg/kg) until day 29, and on day 30 LPS, but not PNU-282,987, was administered. In contrast, a separate cohort of non-lesioned mice (PBS-injected) received asingle dose of PNU on day 30 just prior to the i.p. challenge with LPS (Fig 7A). As shown earlier (Fig 3E), *Sall1* expression in microglia isolated from lesioned animals was significantly lower than from normal microglia (p<0.01, Bonferroni post-hoc, 1-way ANOVA). Multiple PNU-282,987 treatments of such animals resulted in significant upregulation in microglial *Sall1* transcription (p<0.05, Bonferroni post-hoc, 1-way ANOVA), compared to vehicle-treated lesioned group. We thus analysed the expression of the priming hub genes *Clec7a* and *Itgax*, which we had shown were elevated in microglia from lesioned animals (Fig. 3E). Multiple PNU-282,987 treatments markedly reduced the levels of Clec*7a* and *Itgax* (both p<0.001, Bonferroni post-hoc, 1-way ANOVA), bringing them close to baseline levels (i.e. unlesioned PBS controls), although in the case of *Clec7a* these remained somewhat elevated with respect to controls (Fig. 7B-D). Microglial *Il1b* was very robustly increased in p75-saporin+LPS animals (p<0.01) but multiple PNU-282,987 treatments significantly suppressed this microglial *Il1b* response even though it was administered distal to the time of LPS treatment (p<0.001, Bonferroni post-hoc, 1-way ANOVA). Therefore multiple consecutive nicotinic agonist treatments restore the homeostatic gene *Sall1*, suppress the priming genes *Itgax* and *Clec7a* and, therefore, subsequent responses to systemic LPS challenge were normalised (Fig. 7E). Conversely, treatment with PNU-282,987 had no significant effects on these parameters in non-lesioned animals. This suggests that in animals with normal cholinergic tone, and a homeostatic microglial state (based on high *Sall1* and low *Itgax* and *Clec7a*), the nicotinic agonist PNU-282,987 does not impact on the microglial response to LPS. Flow cytometric analysis of single cell suspensions obtained from hippocampi of p75-saporin (lesioned) and PBS injected (unlesioned) animals also revealed a significant (p<0.001, 4 fold approx.) increase in CD45^low^CD11b^+^ microglia frequency in lesioned animals injected with saline or LPS when compared to PBS-injected animals, and this was also mitigated in PNU282,987-treated mice (p<0.01, Bonferroni post hoc, 1-way ANOVA) (Fig. 7F-G).

**Figure 7:**
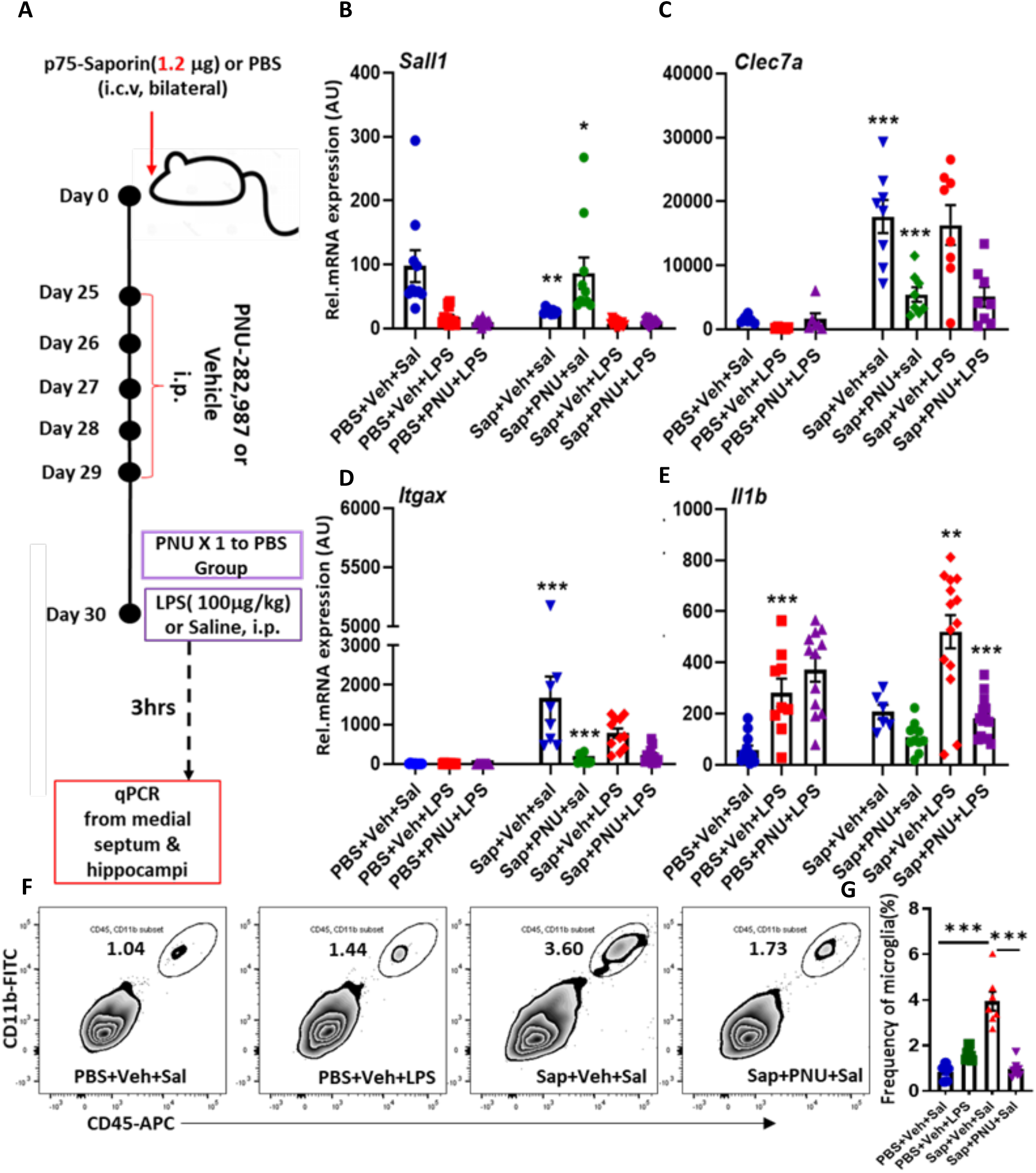
Multiple PNU-282,987 treatments attenuates microglial priming and LPS-induced Il1b. (A) Schematic showing timeline of multiple (5X, 10mg/kg) PNU-282,987 administration, LPS injection (250 µg/kg) and tissue sampling for FACS isolation in lesioned (1.2µg) and PBS control mice. PBS control mice received only single dose (10mg/kg) of PNU-282,987 (A). Graphs showing expressions of Sall1 (marker of resting state) (B), Clec7a (C) and Itgax (D) in isolated microglia from lesioned and PBS injected mice receiving either LPS or saline (sal) and PNU-282,987 or veh. Graphs showing expressions of *Il1b* in isolated microglia from lesioned and PBS injected mice receiving either LPS or saline (sal) and PNU-282,987 or veh (E). Representative flow cytometry dot plots showing effect of 5X PNU treatment on microglia (CD45low CD11b+) numbers in lesioned (1.2µg) and PBS control mice, receiving LPS or saline (sal). Analysed by ordinary One-way ANOVA, significant differences are denoted *p < 0.05, **p < 0.01, and ***p < 0.001 by Bonferroni post-hoc analysis.

## DISCUSSION

Here we have shown that the withdrawal of basal forebrain cholinergic tone from the hippocampus, through p75-saporin lesioning of the medial septum, causes microglia to diverge from their homeostatic state and become primed. In this primed state they are hypersensitive to subsequent peripheral inflammatory stimulation with LPS, producing exaggerated levels of IL-1β. Using novel biosensor technology we show that LPS treatment acutely increases the release of ACh in both the hippocampus and frontal cortex, which was prevented by prior basal forebrain lesions. We demonstrate α7 receptor expression on isolated microglia and show that the primed phenotypic state was reversed by the activation of nicotinic receptors using the α7 agonist PNU-282,987, and this facilitated the suppression of exaggerated IL-1β responses to systemic LPS. The data indicate that basal forebrain ACh is important in maintaining the homeostatic microglial state and that loss of cholinergic tone leads to priming of microglia and hypersensitive neuroinflammatory responses to secondary inflammatory stimuli.

### Loss of cholinergic tone leads to persistent microglial priming

A gradual but significant loss of cholinergic innervation occurs in the ageing brain and particularly in those with cognitive decline and AD (Schliebs & Arendt, 2011) (Kerbler *et al*., 2015). It has emerged that microglial-mediated neuroinflammation is a contributor to the development of Alzheimer’s disease, but the relationship between loss of cholinergic tone and microglial activation has received limited attention. The major contributor to forebrain ACh are the clusters of cholinergic neurons located in the basal forebrain, which project to neocortex, hippocampus and amygdala (Mesulam *et al*., 1983; Woolf, 1991) and here, using the mu-p75-saporin immunotoxin, ChAT neurons in MS and vDBB were selectively targeted while leaving cholinergic cells in the striatum unaffected. The medial septum (MS) and ventral diagonal band of Broca (vDBB) project to the hippocampus via the septohippocampal pathway (Knox & Keller, 2016). The hippocampal CA regions and dentate gyrus are innervated by fibres emanating from MS in a laminar fashion: fibres originating from the vDBB project to CA1 pyramidal and dentate granule cell layers in the dorsal hippocampus, whereas those from the mediolateral DBB send inputs to all CA fields in the caudal hippocampus via the medial part of the fimbria (Teles-Grilo Ruivo & Mellor, 2013). Using ChAT IHC and qPCR we observed substantial but quite variable loss of cholinergic neurons in MS and vDBB. Therefore this required a method to assess septal ChAT depletion in each cellular/molecular experiment rather than an initial quantification of cholinergic neuronal loss in neuropathological analyses. We have previously shown that qPCR for *Chat* in the MS provides a good proxy for ChAT-positive neuronal cell loss in the septum Bonferroni post-hoc, 1-way ANOVA) and we therefore performed correlations of hippocampal microglial gene expression with septal *Chat* in the same animal, showing approximately 50-70% decreases in *Chat* with 1.2 µg of p75^NTR^-saporin roughly consistent other studies using this toxin (Hunter *et al*., 2004; Moreau *et al*., 2008), although some prior studies did show almost complete loss of septal ChAT-positive neurons with this dose (Nag *et al*., 2009).

The cholinergic loss in MS and vDBB gave rise to robust loss of hippocampal ChAT-positive terminals and this was associated with microglial activation, showing increased numbers, altered morphology and persistent activation. The expression of pro-inflammatory cytokines occurred rapidly upon lesion but reduced substantially after 7 days. However microglial markers associated with phagocytic activation, such as *Cd68* and *Trem2* rose more slowly and remained unresolved for at least 35-40 days, similar to microglial activation in neurodegenerative conditions (Schwartz & Baruch, 2014). Though these cells remain morphologically activated it is only upon further stimulation that they robustly produce IL-1β, akin to the original descriptions of microglial priming in neurodegeneration (Cunningham *et al*., 2005) nd ageing (Godbout *et al*., 2005). The persisting microglial activation was also characterised by loss of the homeostatic state regulator *Sall1* and the increased expression of the previously described priming hub genes *Itgax, Clec7a* and *Axl* (Holtman *et al*., 2015) and *C3, C1q, Trem2* and *Tyrobp* which are strongly associated with neurodegeneration and synaptic loss (Hong *et al*., 2016; Keren-Shaul *et al*., 2017). Significantly, in the case of each ‘priming’ transcript, changes in hippocampal expression of these genes was strongly correlated with septal *Chat* levels: hippocampal *Sall1* was directly correlated with septal *Chat* while *Itgax, Clec7a* and *Axl* were all indirectly correlated with septal *Chat*. Those relationships indicate that, even at a time point distal to the initial cholinergic lesions, the microglia lose their homeostatic signature and remain in a primed state that is proportionate to the extent of loss of cholinergic tone. The propensity of these microglia to produce exaggerated IL-1β responses to LPS-induced acute systemic inflammation was also directly correlated with Cd11c expression, the product of *Itgax*, therefore it is clear that the magnitude of expression of transcripts of the primed signature predicts the extent of exaggerated IL-1 responses.

### Relationship to cholinergic signalling

These data are consistent with prior demonstrations that ACh exerts anti-inflammatory influence on peripheral macrophages, suppressing LPS-induced IL-1β and TNF-α expression (Wang *et al*., 2003) agonism at the α_7_-nAChR (Tracey, 2007). There are several studies indicating that nicotinic agonism also reduces neuroinflammation (see introduction). Stimulation of the α_7_-nAChR increases the expression of anti-inflammatory cytokines TGFβ1, IL-4 and IL-10 in cultured microglia (De Simone *et al*., 2005; Rock *et al*., 2008; Parada *et al*., 2013; Zhang *et al*., 2017), but detailed *in vivo* studies researching the nature of ACh and α_7_-nAChR control of microglial phenotype have been lacking. The current data clearly show that isolated microglia express α_7_AChR, that withdrawal of cholinergic tone persistently alters microglial phenotype and that nicotinic agonism, with PNU-282,987 reduced the levels of IL-1β induced by systemic LPS. Although prior studies with LPS-induced sepsis have shown that nicotinic agonists and/or electrical stimulation of vagal ACh outflow reduces LPS-induced circulating cytokines (Wang, Yu et al. 2003), IL-1β, IL-6 and TNFα were largely unaffected by the low dose of PNU-282,987 (10mg/kg) in the current study, indicating that PNU treatment-mediated downregulation of LPS-induced IL-1β in lesioned hippocampus is independent of effects on peripheral inflammation. The much lower doses of LPS used here may be significant. We tentatively suggest that the peripheral cholinergic anti-inflammatory pathway may restrict the severity of profound inflammatory challenges, such as severe sepsis, but does not interfere with more adaptive inflammatory responses to more moderate inflammatory stimuli (100µg/kg LPS). When we used a higher PNU-282,987 dose (20mg/kg) this did appear to lower expression of circulating cytokines (TNF-α, IL-1β).

As previously discussed ACh signalling may be significantly reduced in the brains of those with neurodegenerative disease such as AD and this has been proposed to be important for disease-associated changes via the α7-nAChR. This receptor has been suggested to be neuroprotective in multiple models of brain pathology but direct effects of nicotine are observed on neurons and astrocytes as well as on peripheral macrophages and microglia (Hijioka *et al*., 2012; Liu *et al*., 2012b; Han *et al*., 2014; Guan *et al*., 2015). We aimed to clarify the ways in which ACh might exert its anti-inflammatory influence using the current model systems. Firstly, using an implantable biosensor to measure choline in real-time (as a proxy for ACh) (Baker *et al*., 2015; Teles-Grilo Ruivo *et al*., 2017; Baker *et al*., 2018) we found that acute LPS treatment triggered an increase in cholinergic tone with an onset of <5 minutes, but which lasted for a number of hours, indicating relatively rapid, though clearly not phasic, release of ACh. That this was prevented in lesioned animals indicates that basal forebrain neurons are responsible for this release. This could, in theory, have anti-inflammatory functions, immediately exerting a modulating influence on LPS-induced inflammation. Nicotine has been shown to reduce LPS-induced circulating cytokines, consistent with the cholinergic anti-inflammatory pathway exerting its effects in the periphery (Tracey, 2007), but this was with significantly higher LPS dosing (Kojima *et al*., 2011). In our experiments the nicotinic agonist PNU-282,987 did not have any significant effect on LPS-induced hippocampal cytokines in non-lesioned animals (figures 5,7). This indicates that in normal animals acute changes in nicotinic signalling have limited ability to suppress LPS-induced acute neuroinflammatory changes. Rapidly induced ACh may, more plausibly, be important in neurovascular responses to systemic changes arising from acute LPS, since acutely enhanced ACh tone is known to potentiate cerebral blood flow and facilitate potentiated neuronal responses (Lecrux *et al*., 2017). The simultaneous increases in hippocampus and frontal cortex are also consistent with coordinated hippocampal and frontal cortex release in animals showing higher arousal during maze performance (Teles-Grilo Ruivo *et al*., 2017). How this is related to the low arousal state that begins in the hours after LPS is not clear but the tonic increase in choline does appear to begin to resolve from approximately 3 hours. ACh was previously shown, by microdialysis, to be briefly elevated after intracerebroventricular IL-1β administration before showing suppressed levels for a number of hours at least (Taepavarapruk & Song, 2010). The current study uses higher temporal resolution and early events indicate an initially increased cholinergic activity in the hippocampus followed by a slow decay towards baseline levels. Both AChE and α7-nAChR have been reported to be decreased in the days after LPS-induced sepsis (Lykhmus *et al*., 2016), suggesting that ACh or nicotinic agonists may lose their anti-inflammatory potency after stimulation. Since the cholinergic lesions used here ablate the acute ACh increases this would likely hamper any such neurovascular or indeed neuromodulatory actions.

Given the effects observed on responses to LPS in lesioned animals we instead proposed that persistent nicotinic agonism might mimic the effects of ACh in the hippocampus and restore the normal homeostatic microglial profile. The daily treatment of lesioned animals with PNU-282,987, for 5 days, but without treatment on the day of LPS challenge, significantly increased the expression of the homeostatic gene *Sall1*, reduced the expression of priming genes *Clec7a* and *Itgax* and prevented the exaggerated synthesis of IL-1β upon LPS challenge (figure 6). This tonic nicotinic agonism thus returns microglia towards their homeostatic state. Some limited expression of *Clec7a* and *Itgax* persists after PNU-282,987 treatments and these may be explained by cellular debris and process degeneration that remain even 40 days after the degenerative effects of the lesion. Bolstering cholinergic signalling is unlikely to be able to reverse priming/activation by all stimuli, but is clearly sufficient to ‘reset’ at least those primed by loss of ACh. Flow-cytometric analysis revealed that restoration of cholinergic tone in hippocampus of lesioned animals was also successful in reducing lesion-induced microglial proliferation. This is consistent with previous studies where PNU treatment(s) reduced microglial numbers observed in animal models of PD and migraine (Stuckenholz *et al*., 2013; Liu *et al*., 2018).

The signalling mechanisms by which α7 cholinergic activity restores the microglial homeostatic phenotype have not been investigated here. There is not clear evidence that nicotine induces membrane currents in microglia, but there is some evidence that α7 receptors can also signal in a metabotropic mode (Kabbani & Nichols, 2018). The anti-inflammatory effect of nicotine on monocytes and macrophages appears to involve induction of Jak2/STAT3, induction of PI3K, inhibition of p38 MAP kinase and inhibition of NFkB, signalling, contributing to suppression of TLR signalling and release of mediators such as TNF-α (Kalkman & Feuerbach, 2016). However those prior studies largely related to inhibition of LPS-induced pathways, which, in microglia, are clearly distinct from those that characterise the ‘primed’ phenotype (Holtman *et al*., 2015). Conversely, TGFβ1 is a major regulator of the homeostatic microglial phenotype (Butovsky *et al*., 2014; Krasemann *et al*., 2017) and this has been shown to be induced by nicotine (Rock *et al*., 2008) although this remains little studied. *Sall1* is a transcription factor that acts as a key regulator to retain the homeostatic phenotype (Buttgereit *et al*., 2016). In that study conditional knockout of TGFβ1 also suppresses *Sall1* expression shifting microglia to an activated phenotype. It will be important to investigate the impact of loss of nicotinic cholinergic tone on the interaction of these factors *in vivo*.

### Implications of loss of cholinergic tone in dementia, sepsis and delirium

Much has already been written about potential therapeutic applications of nicotinic signalling for brain disease. Preclinical studies have shown beneficial effects of α7-nAChR agonists in neurodegenerative diseases such as Parkinson’s disease (Stuckenholz *et al*., 2013) (Liu *et al*., 2012a), Huntington’s disease (Foucault-Fruchard *et al*., 2018) and in Alzheimer’s disease (Medeiros *et al*., 2014) (Vicens *et al*., 2017). However the demonstration, here, of the impact of basal cholinergic tone on the homeostatic phenotype of microglia helps to clarify the nature of anti-inflammatory effects of nicotinic agonists but also shows that the loss of cholinergic tone, in aging and early dementia, likely reduces the brain’s resilience to secondary inflammatory challenges such as infection, surgery or injury. Such acute insults occurring in the context of aging or dementia have multiple adverse outcomes for patients, including delirium and accelerated progression of neurodegeneration in humans (Fong *et al*., 2009; Witlox *et al*., 2010; Wilson *et al*., 2020) and experimental animals (Cunningham *et al*., 2009; Field *et al*., 2012; Skelly *et al*., 2019). We have previously showed that p75-saporin lesions left the brain vulnerable to acute and transient working memory deficits induced by LPS independent of microglial priming (Field *et al*., 2012). The current study used more robust cholinergic lesions to now also leave microglia primed and it will be important to assess the cognitive consequences of this increased brain vulnerability. Given the utility of nicotinic agonists as anti-inflammatory agents in preclinical models of sepsis (Tracey, 2007) and the acute and long-term cognitive sequelae of sepsis/critical illness (Iwashyna *et al*., 2010; Widmann & Heneka, 2014; Annane & Sharshar, 2015), it was tempting to speculate that bolstering cholinergic function might reduce delirium in patients with sepsis, but the cholinesterase inhibitor rivastigmine was shown not to reduce the incidence of delirium in critically ill patients (van Eijk *et al*., 2010). Given the neuromodulatory and immunomodulatory actions of nicotinic agonists, one might predict that these agonists might provide a more successful approach, but it also may be that such drugs would be most promising in patients that have existing cholinergic degeneration since the restoration of cholinergic tone had very substantial effects on the brain response to acute inflammatory stimulation in the current study. Likewise there is some preclinical evidence that sepsis leads to damage within the cholinergic system (Semmler *et al*., 2007; Silverman *et al*., 2015; Zaghloul *et al*., 2017), which may further increase the vulnerability of the brain to subsequent inflammatory insults.

## Conclusion

The data presented here illustrate that loss of cholinergic tone in the hippocampus, arising from the medial septum, drives microglial activation in the hippocampus. Microglia lose their homeostatic status and become primed resulting in hyper-sensitivity to subsequent inflammatory insults. It will be important to elucidate the consequences of this loss of cholinergic tone for cognitive and neuropathological outcomes in neurodegeneration, sepsis and delirium.

